# Continuous interdomain orientation distributions reveal components of binding thermodynamics

**DOI:** 10.1101/200238

**Authors:** Yang Qi, Jeffrey W. Martin, Adam W. Barb, François Thélot, Anthony Yan, Bruce R. Donald, Terrence G. Oas

## Abstract

The flexibility of biological macromolecules is an important structural determinant of function. Unfortunately, the correlations between different motional modes are poorly captured by discrete ensemble representations. Here, we present new ways to both represent and visualize correlated interdomain motions. Interdomain motions are determined directly from residual dipolar couplings (RDCs), represented as a continuous conformational distribution, and visualized using the disk-on-sphere (DoS) representation. Using the DoS representation, features of interdomain motions, including correlations, are intuitively visualized. The representation works especially well for multidomain systems with broad conformational distributions. This analysis also can be extended to multiple probability density modes, using a Bingham mixture model. We use this new paradigm to study the interdomain motions of staphylococcal protein A, which is a key virulence factor contributing to the pathogenicity of *S. aureus*. We capture the smooth transitions between important states and demonstrate the utility of continuous distribution functions for computing components of binding thermodynamics. Such insights allow the dissection the dynamic structural components of functionally important intermolecular interactions.

## 3 Abbreviations

see footnote^1^.

## 4 Introduction

Dynamics of biological macromolecules over various spatial and time scales are essential for biological functions such as molecular recognition, catalysis, and signaling. Multiple studies have demonstrated that interdomain motions are a crucial component of the dynamics associated with molecular recognition [1, 2, 3]. Protein and RNA molecules can rearrange interdomain orientations upon binding, thereby enabling adaptive conformational changes that are accompanied by contributions to the free energy, internal energy, and entropy of binding. Despite the recent interest in studying interdomain motions, one challenge of studying such motions is the lack of effective representations. Using the discrete ensemble representation, correlations between different motional modes are not directly captured. Here, we present a new paradigm to represent and visualize correlated interdomain motions. We represent interdomain motions as a continuous distribution on *SO*(3) (3D rotational space) and parameterize the correlations directly. By using the continuous representation, correlations are quantified directly from experimental observables. In addition, this continuous representation enables the computation of the re-orientational components of the free energy, internal energy, and entropy of binding. We also design a visualization method called the disk-on-sphere (DoS) drawing to visualize the spatial dynamic information (see Figure 1). Features of the interdomain motions, including geometry and correlations of these motions, are effectively represented in the DoS visualization. Finally, unlike discrete ensemble representations, the continuous representation of orientational distributions is not over-fit, requires no non-physical assumptions, and can be computed directly from the measured RDCs (see section 3.4 below).

**Figure 1:**
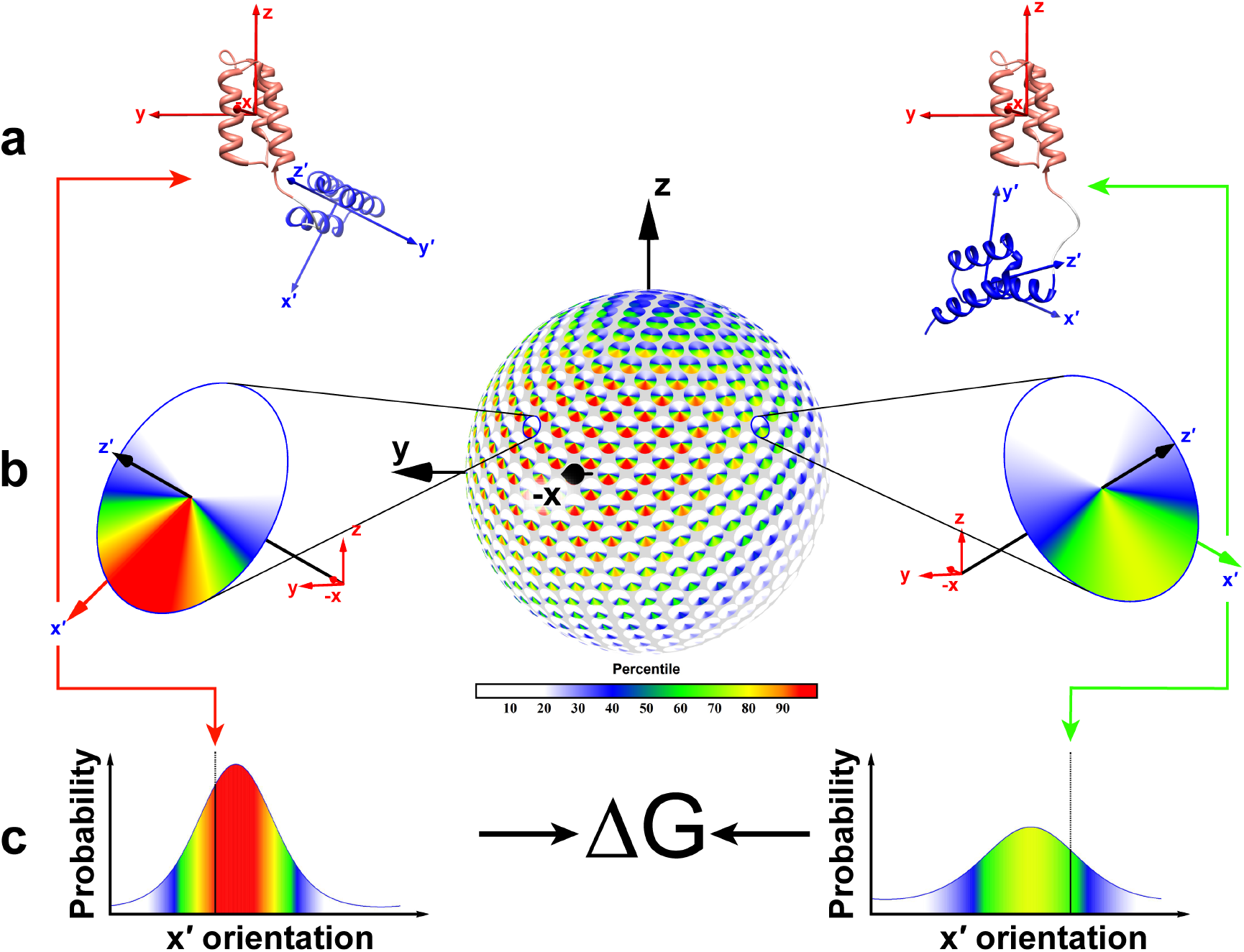
Summary of continuous distribution of interdomain orientations (CDIO) analysis and the Disk-on-Sphere (DoS) representation. Interdomain orientations are defined as the orientation of the unaligned domain (blue, *x′*,*y′*,*z′*) in the aligned domain’s coordinate system (red, *x*,*y*,*z*). In the ZLBT-C system, the aligned domain is the ZLBT domain and the unaligned domain is the C domain. **a**, Two examples of interdomain orientations. The probabilities of each of these orientations is deduced from the best-fit CDIO determined from observed residual dipolar couplings in the unaligned domain. **b**, The DoS plot of one of two best-fit CDIO solutions. The position of each disk on the sphere gives the orientation of the *z′* axis in the *x*,*y*,*z* frame. The colors on each disk represent the joint probability of a particular orientation of the *x′* and *z′* axes. The two enlarged disks correspond to the interdomain orientations depicted in panel **a**. **c**, The conditional probability distributions of each of the interdomain orientations in panel **b**. The ratio of the probabilities of the two orientations can be used to deduce their difference in Gibbs energy.

Staphylococcal protein A (SpA) is a key virulence factor that supports the invasion of *Staphylococcus aureus* into the human body. SpA binds to an array of targets in the host to disarm the immune system, facilitate the colonization and consequently contribute to the pathogenicity of *S. aureus*. The emergence of antibiotic-resistant *S. aureus* strains has driven the search for vaccines with high efficacy to counteract several virulence factors, including SpA [6]. Targeting virulence as opposed to bacterial cell growth may be a more effective way to avoid the emergence of resistant strains. Structural studies of SpA and its interaction with host proteins would support rational design of improved vaccines or other therapeutics to diminish *S. aureus* virulence. The binding targets of SpA include the F_c_ region of antibodies, the F_ab_ region of V_H_3 antigen receptors (e.g., IgM) on B cells, TNFR1, and EGFR [7, 8, 9]. There are five tandem functional domains in the N-terminal half of SpA (SpA-N). The five domains share a high sequence identity and they are structurally and functionally similar [10, 11, 8]. Recent studies indicate a correlation between the functional plasticity and the structural flexibility of SpA-N [10, 13, 12].

Given the limited current knowledge of SpA-N interdomain motions, we undertook an NMR study as an initial effort to understand the link between structural flexibility and functional plasticity. NMR spin relaxation experiments identified a 6-residue flexible interdomain linker and interdomain motions. To obtain spatial dynamic information, we measured residual dipolar couplings (RDCs) from two SpA-N domains with multiple alignments. The interdomain motions of SpA-N were analyzed using the new paradigm. Significant correlations were observed in the orientational distribution, indicating that the motion populates some interdomain orientations more than others. A novel statistical thermodynamic analysis of the observed orientational distribution suggests that it is among the energetically most favorable orientational distributions for binding to antibodies. Thus, the affinity is enhanced by a pre-posed distribution of interdomain orientations while maintaining the flexibility is presumably required for function.

## 5 Results

### 5.1 The linkers between SpA-N tandem domains are highly flexible

SpA-N has five tandem domains (E-D-A-B-C). Each domain is a three-helix bundle. NMR structural studies suggest that there are unstructured regions between domains, each consisting of 6 to 10 residues [14]. However, the flexibility of these unstructured regions has not been directly measured. Hence, we conducted NMR spin relaxation experiments to measure the flexibility at picosecond to nanosecond timescale. Instead of using the five domains of SpA-N, we replaced E/D/A/C domains with B domain and engineered a construct called 5B [13]. Because the sequence identity between domains is high (83 - 91%), this construct is a mimic of SpA-N structurally and functionally. Compared to SpA-N, the construct has a simpler ^15^N-HSQC spectrum. Except for the N- and C-termini, the residues at the same position in each domain have nearly the same electromagnetic environment, so their signals in a ^15^N-HSQC spectrum overlap (Figure S1). The 55 overlapping ^15^N-HSQC resonances corresponding to all 5 domains of 5B were sequentially assigned using standard methods. Six additional resonances were assigned to the first five residues at the N-terminus and the last residue at the C-terminus. Those terminal residues would experience different electromagnetic environments from the corresponding interdomain linker residues because they have no adjacent domain.

We measured R1 and R2 rates and heteronuclear NOE ratios at 14.1 T for each assignable residue using published methods [15]. A sharp drop in the heteronuclear NOE ratios was observed for the first five residues and the last one residue in each domain (Figure S2). We also generated a Lipari-Szabo map by deriving spectral density estimates from the relaxation experiments (Figure 2a). The Lipari-Szabo mapping method provides a graphical approach to estimate and display order parameters [16]. The majority of points in Figure 2a form a cluster near the bottom of rigid limit curve. For the corresponding residues, reorientation of the backbone N-H bond is primarily due to global tumbling. However, 9 points are well separated from the cluster, indicating that the N-H bonds of these residues are much more flexible. Four of these points correspond to residues 3-5 and the C-terminal residue of 5B and are assigned to the termini because they have the lowest order parameter for the sequentially-assigned residue. Their low order parameters are typical for the termini of proteins. The remaining low order parameter N-H bonds correspond to residues 1-5 and 58 of each non-terminal B domain. The order parameters of these N-H bonds are nearly as low as those of the termini. The heteronuclear NOE ratios (Figure S2) and the Lipari-Szabo mapping (Figure 2a) are consistent with each other. Both of them show that there is a six-residue linker (KADNKF) between every pair of adjacent domains. While linkers between D-A-B-C domains of SpA-N have the same sequence as the linker in 5B, the linker between E and D domains has three additional residues. As a result, the E-D domain linker could be more flexible than others. The motions of each domain could deviate from isotropic tumbling because the motions are also affected by the motions of adjacent domains. Indeed, the Lipari-Szabo mapping plot does not have the appearance of one generated from a globular protein. Using an anisotropic tumbling model, we estimated the three global correlation times as 11.02, 11.91 and 14.58 ns, which are much smaller than the correlation time of a globular protein of the size of SpA-N. The small correlation time implies that there are interdomain motions due to the flexible linkers.

**Figure 2:**
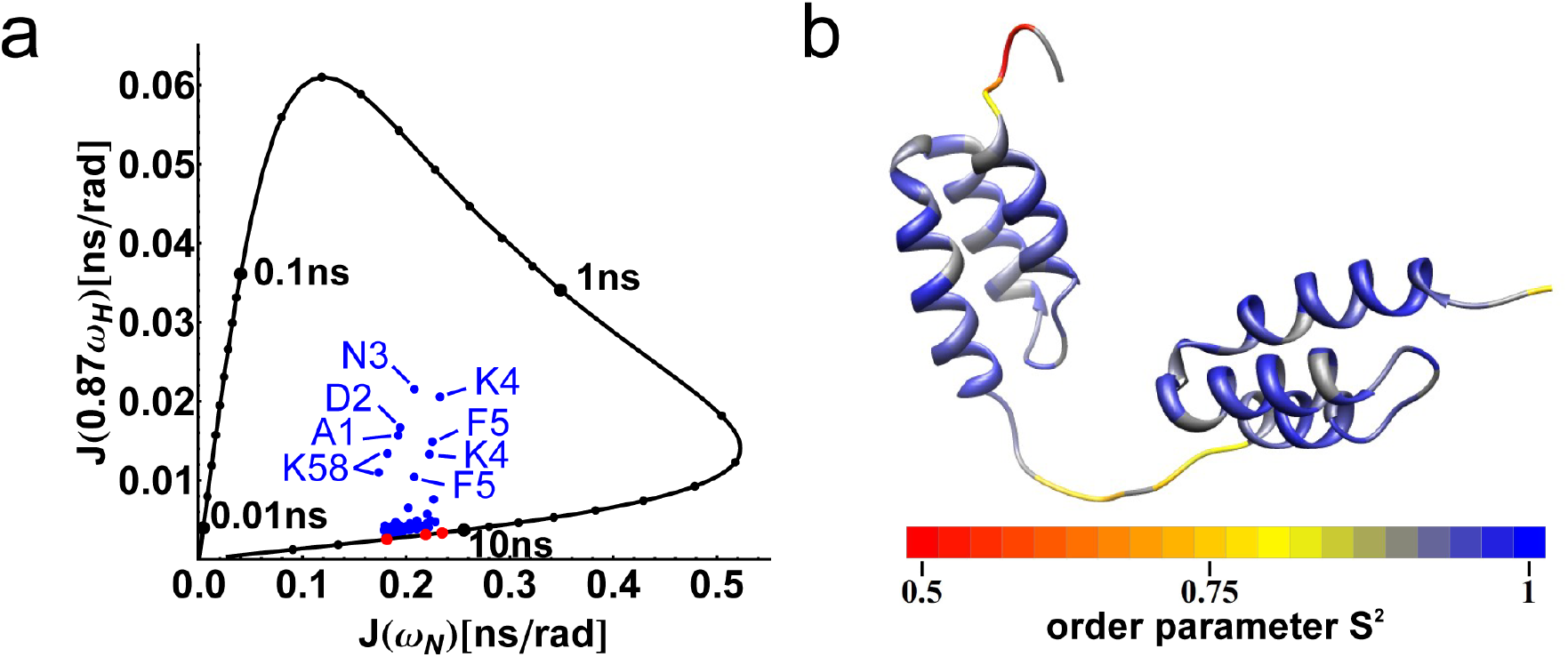
Relaxation properties of 5B. **a**, Lipari-Szabo mapping of 5B. Residue resonances are plotted as blue points inside the rigid limit curve. Anisotropic correlation times are plotted as red points on the rigid limit curve. **b**, Order parameters are color coded onto a structure of double B domains. Residues are colored as gray when data is not available.

### 5.2 Partial alignment of a reference domain decouples alignment from interdomain motion

We measured residual dipolar couplings (RDCs) of a di-domain construct that links a B domain variant “Z domain” [17] and a C domain together using the conserved linker. Z domain differs from B domain by only two residues, A1V and G29A. Consequently, they have nearly indistinguishable structures [17, 10]. RDCs are sensitive to motion from picoseconds to milliseconds, so our measured RDCs are averaged over local bond vibration, interdomain motion, and global tumbling. Based on the results from spin relaxation experiments, residues outside of the linker regions are rigid and the contribution from local bond motions is negligible. In order to separate interdomain motions from global motions, we introduced a rigid lanthanide binding tag (LBT) [18] into the loop between Helix II and Helix III of Z domain to create the ZLBT domain [19] in the N-terminal domain, linked to C domain at the C-terminus (referred to hereafter as ZLBT-C). This construct allowed us to observe the interdomain motions in the Z (B) domain coordinate system. Thus, the ZLBT domain is directly aligned by a lanthanide ion while the C domain is indirectly aligned. The alignment of the C domain depends on both global tumbling and interdomain motions. Because the alignment of the ZLBT domain only affects its global tumbling, we could remove the effect of global tumbling from the alignment of the C domain and consequently decouple interdomain motions from global motions.

In order to obtain multiple alignments, we constructed a second version of lanthanide-binding ZLBT domain (NHis-ZLBT-C) by introducing a His-tag at its N-terminus (Figure S3). Both constructs have nanomolar lanthanide binding affinities as measured by Tb^3+^ fluorescence (Figure S3). The construct NHis-ZLBT-C has a tighter binding affinity, suggesting that the His-tag contributes to the coordination of lanthanide ion. Because the His-tag is in close proximity to the LBT, the tighter binding affinity could be due to a direct interaction between the His-tag and the lanthanide ion. Alternatively, the His-tag may favor the higher affinity conformation of LBT in some other way, indirectly enhancing the binding of the lanthanide ion to LBT. Either way, the introduced His-tag could perturb the detailed conformation of the LBT-lanthanide complex and potentially result in a different alignment. Indeed, alignments from the two constructs are different (Table S1). We also combined the two constructs with two types of lanthanide ions, Dy^3+^ and Tb^3+^. The two lanthanides have different magnetic susceptibility tensors and behave differently in a magnetic field, so the combination gave us a total of four different alignments (Table S1). The RDCs in all four alignments were measured by ^15^N IPAP HSQC experiments (see Materials and Methods). The RDCs of C domain in all four alignments are significantly smaller than those of B domain, indicating extensive interdomain motions (Figure S5). The small RDCs of C domain were carefully measured by fitting the peaks to a mixture model with both Gaussian and Lorentzian components as described in Materials and Methods. We performed duplicate experiments to estimate the RDC error. The estimated error is 0.2 Hz, which agrees with literature estimates and the RMSDs in the correlation plots (Figure S5) [20]. Because no significant intra-domain dynamics were observed within the picosecond to nanosecond timescale, our analysis treats the non-linker region of each domain as a rigid body. We fit Saupe alignment tensors to the RDCs from the non-linker residues of each domain in each alignment [21]. There is a good agreement between experimental data and back-calculated data (Figure S5), which in turn confirms the validity of the rigid body assumption for the individual domains. Fit results are summarized in Table S1.

### 5.3 The information content of the RDC data can be determined using the OLC method

For interdomain motions, the Saupe tensors summarize the information content in the RDC data sets [2]. We quantified the information content in the four sets of RDC data using the orthogonal linear combination (OLC) method [22]. The method projects different alignments in the five dimensional Saupe tensor space and finds a linear combination of these to yield a set of orthogonal alignment tensors. The information content of these orthogonal alignments can be quantified from the Q factors for each alignment. The number of orthogonal alignments with low Q factors correlates with the constraining power of the data. By quantifying the information content, we could build a model matching the constraining power of our data and avoid over-fitting. In addition, some OLC components contain a considerable amount of noise giving them low information content. Filtering these components out can reduce noise in the derived orientations.

Out of four different alignments, we obtained two orthogonal alignments with high information content (Figure 4a). The singular value measures the amplitude of signal in the corresponding alignment while the Q factor measures the signal-to-noise ratio. The first orthogonal alignment has a large singular value of 34.4 × 10^−4^ and a low Q factor of 0.13, indicating high information content. Although the second orthogonal alignment has a much smaller singular value, its Q factor of 0.22 is still below 0.3, suggesting a much smaller noise and high information content. The third and fourth alignments have low singular values and high Q factors, so they have low information content. Consequently, the first two alignments were used for constraining the conformational model while the rest were discarded. With the two orthogonal alignments (Table S2), we have a total of 10 independent observables to constrain a model of the interdomain orientational distribution.

### 5.4 Interdomain motions can be represented as a continuous distribution

Previous approaches traditionally use a discrete finite ensemble to describe biomolecular dynamics, including interdomain motions [1, 2, 3]. A discrete ensemble assigns certain probabilities to the conformations in the ensemble and assigns zero probability to the rest of the conformational space. Although the ensemble description captures several characteristics of biomolecular motions, the void of probability between structures is physically unreasonable. In addition, discrete conformational ensembles in previous studies were only constrained by averages observed in experiments. There is no direct evidence for the atomistic details shown in the discrete conformations. The atomistic details in discrete conformational ensembles are an over-interpretation of the experimental observables. As an alternative to this approach, we took advantage of the insensitivity of RDCs to interdomain distance and modeled the orientational component of the interdomain motions as a continuous distribution on the 3D rotational space *SO*(3). We refer to this henceforth as the *Continuous Distribution of Interdomain Orientations (CDIO)* model. Our CDIO model belongs to the family of Bingham distributions [23] that are widely used to describe circular distributions on the 2D sphere *S*^2^ and 3D rotational space *SO*(3) [24]. Previous studies demonstrated the Bingham model’s ability to represent salient features of a broad spectrum of orientational distributions [24]. Although the Bingham model on *SO*(3) is uni-modal, it is a joint distribution in rotation space that includes all possible interdomain orientations. Unlike discrete ensemble representations, it can accurately describe concerted motions. Fortunately, the Bingham distribution only has nine independent parameters, which can be easily constrained by two or more orthogonal sets of RDC alignments. In contrast to ensemble methods, which rely on a large number of parameters (e.g., molecular mechanics force fields and the size of ensemble), the CDIO model relies only on the assumption that the interdomain orientation is a smooth distribution with a single mode. The smoothness assumption is a weak assumption that excludes high-frequency oscillations in the distribution, thereby preserving the information content in the RDC data, while avoiding over-parameterization.

The CDIO model reduces the structure determination problem to a process of fitting the parameters of the Bingham distribution to the RDC data (see Figure 3). We used a deterministic algorithm to search for the parameters of the distribution. The algorithm belongs to the branch-and-bound family [25, 26, 27], which uses a divide-and-conquer strategy for search problems. The algorithm is also provable, therefore it guarantees to find the best-fit Bingham distribution. To describe the interdomain orientation, it is first necessary to define the molecular frame of each domain. In the coordinate frame of either Z or C domain, the *z*-axis of the domain is parallel to the helices and points toward the N-termini of helices 1 and 3, the *y*-axis is in the plane of helix II and III, and the *x*-axis is perpendicular to the plane (Figure 1a). The 3D joint interdomain orientational distribution can be visualized in a *disk-on-sphere (DoS)* representation (Figure 5). In this representation, the red *x*, *y*, *z*-axes in Figure 1a represent the coordinate frame of the reference domain (Z domain) and the blue *x′*, *y′*, *z′*-axes in Figure 1a represent the coordinate frame of the C domain. The joint probability of a particular interdomain orientation, displayed as a radial line segment representing the *x′* axis of the C domain on a disk whose position on the sphere is determined by the orientation of the *z′* axis, is represented by the color of the line segment. The color scale for the probability, in terms of percentile, is shown by the legend at the bottom of the figure. Note that the line segment color on each disk represents the joint probability of both the *z’* axis orientation and a particular rotation around it.

**Figure 3:**
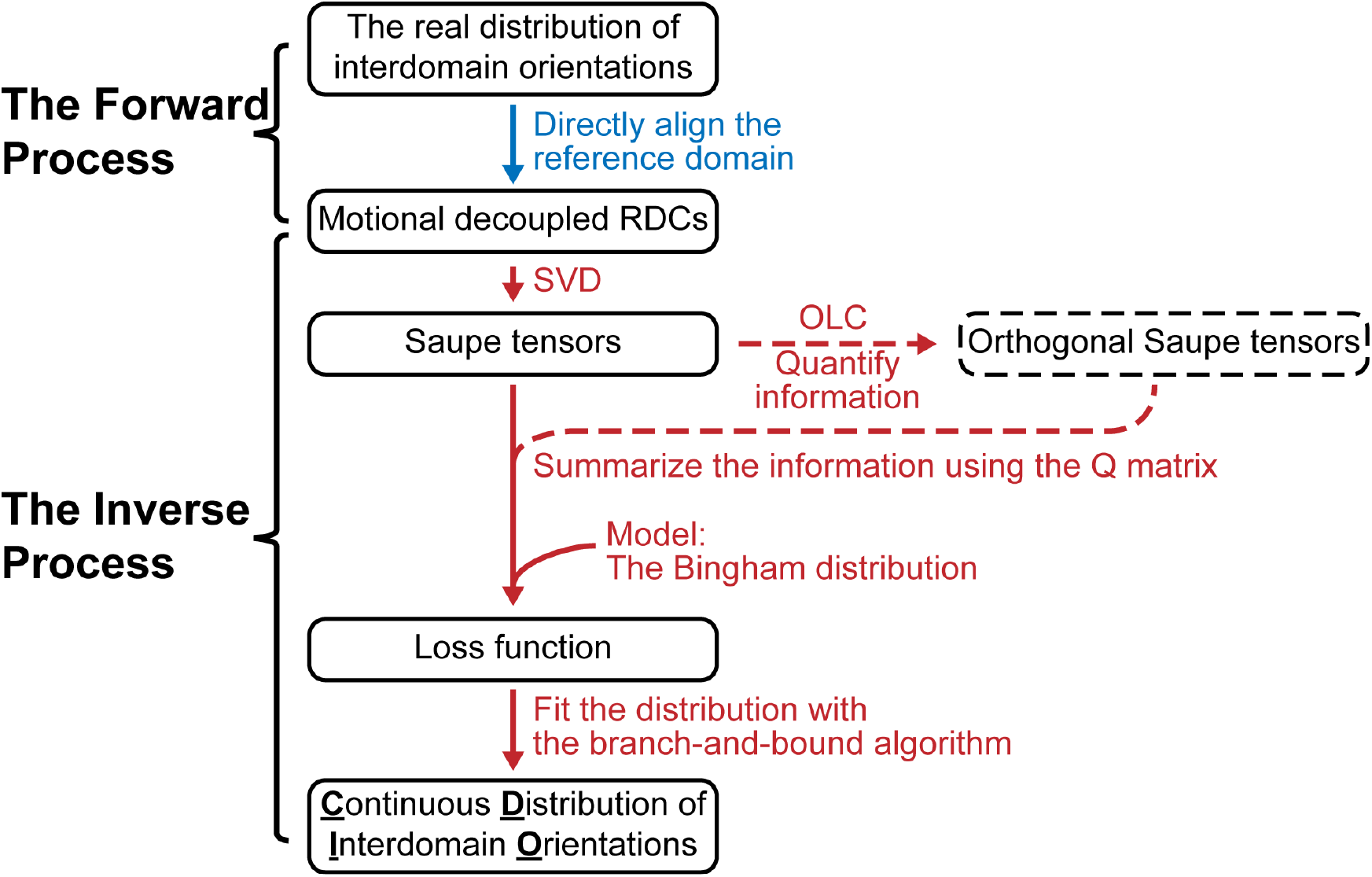
A flow chart of the experimental and computational steps used to calculate the continuous distribution of interdomain orientations. The step in blue is an experimental step. The steps in red are computational steps. The loss function is given in Eq. (33) in SI.

**Figure 4:**
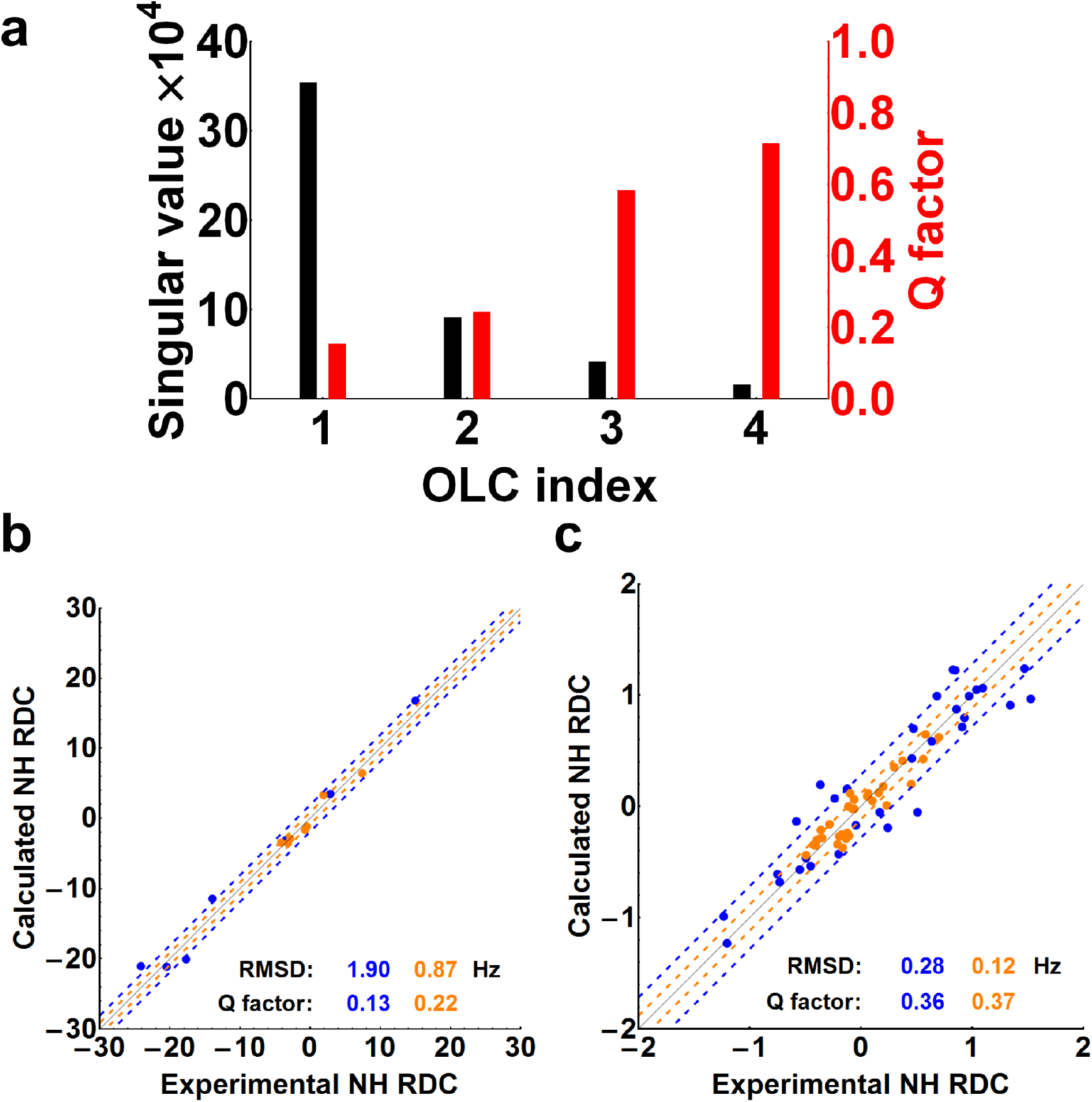
Orthogonal linear combination(OLC) analysis of the four RDC alignments. **a**, Singular values (black bars) and Q factors (red bars) associated with each OLC alignment. **b**, The correlation plot of the first OLC dataset (blue) and the second OLC dataset (yellow) of Z domain and the RDCs back-calculated from the two corresponding OLC Saupe tensors of the Z domain. **c**, The correlation plot of the first OLC dataset (blue) and the second OLC dataset (yellow) of C domain and the RDCs back-calculated from the two corresponding OLC Saupe tensors of the C domain.

Using the two orthogonal alignments generated by the OLC method, we applied the algorithm to the SpA-N di-domain system and found two CDIO model solutions that can reproduce the RDC data nearly equally well (Figure S6). *A priori*, it is possible that any linear combination of the two solutions could also fit the data. To determine if both solutions were physically possible, we simulated a structural ensemble of the B-C di-domain molecule in the absence of any binding partner using the RanCh component of the EOM package [28].

The core Z and C domains were treated as rigid bodies while the backbone conformations of the 6-residue Z-C linker were sampled using RanCh [28] to generate a 10,000 member ensemble. The simulation generated conformers by sampling the Ramachandran space of each flexible residue and rejected conformers with steric clashes. We then converted the resulting discrete ensemble into a CDIO model by convolution of Bingham kernels for each conformer (see Materials and Methods and Figure S8). When comparing the RanCh-generated CDIO (Figure S7) with the experimentally constrained ones, the first solution, shown in Figure 5a, is much less consistent with the simulation than the second (Figure 5b).The DoS plot of the simulation shows a large void of probability in the white area, suggesting that most conformers in this area have steric clashes.

**Figure 5:**
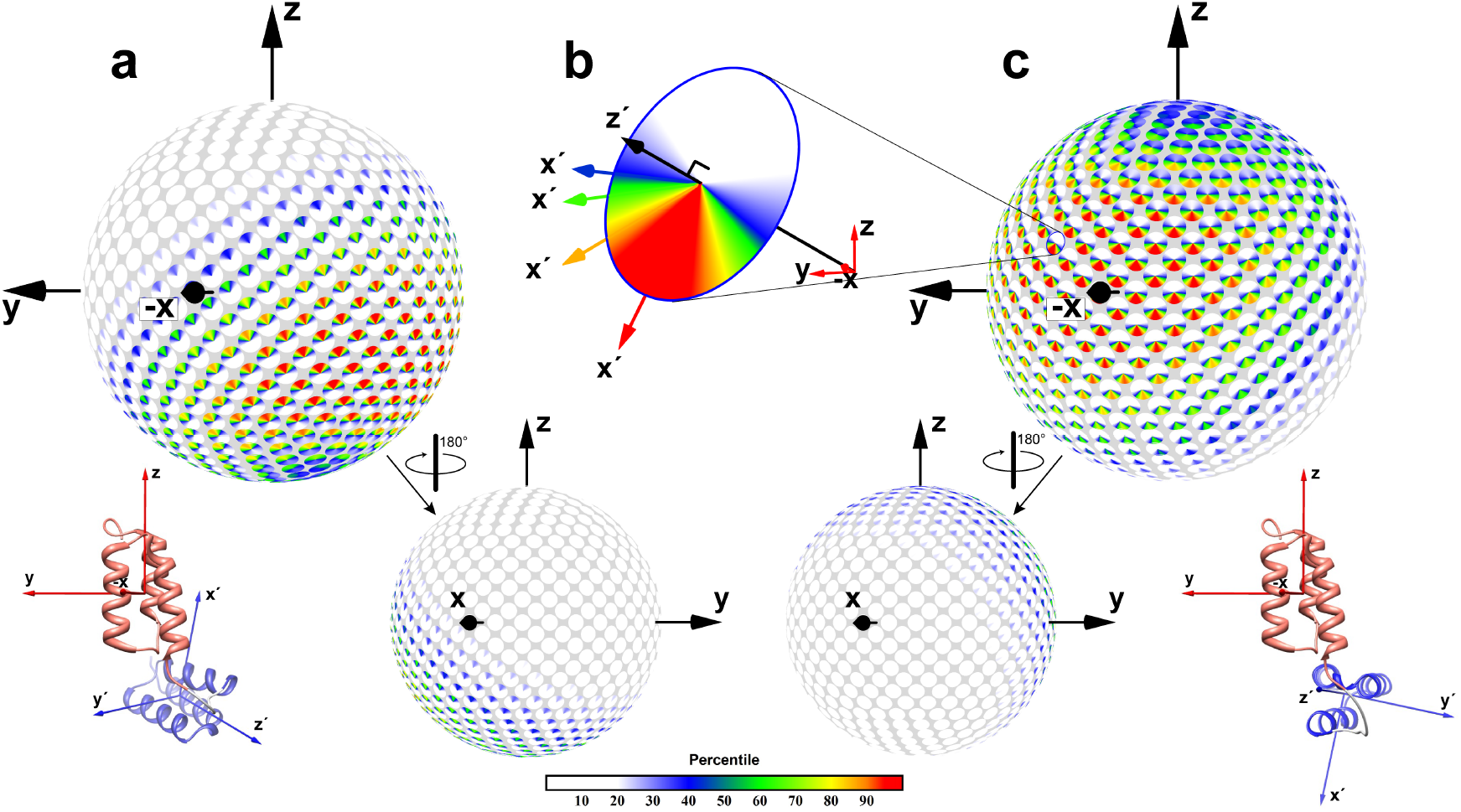
Continuous distribution of interdomain orientations (CDIO) models for ZLBT-C shown in disk-on-sphere (DoS) views. Two solutions give equivalently good fits to the data. The first solution is shown in panel **a** and the second solution is shown in panel **c**. Each panel shows the distribution from a front view (top) and a back view (bottom). Also shown are atomistic models whose interdomain orientation is the most probable in each solution. The linker conformation and interdomain distance are arbitrary. **b**, An example disk showing the joint probabilities of 4 different different interdomain orientations, all with the same *z′* axis orientation but different rotations around *z′*.

The distribution mode of the first solution largely coincides with this low probability region and 37.3% of its probability falls into the region (Figure S7). Quantitatively, we defined a clash score based on the probability of a distribution (see SI) in the low probability region and calculated clash scores for both solutions. The clash score of the first solution is more than two fold higher than that of the second solution. By thermodynamic criteria (see section 5.5 below), the simulated ensemble is also in better agreement with the second solution than the first solution or any linear combination of the two. These results suggest that the first solution is a so-called “ghost” solution [1], but further studies with additional orthogonal alignments are necessary to definitively rule it out.

In the second solution, shown in Figure 5c, the most probable interdomain orientation has the *z′*-axis of C domain close to the minus *x*-axis of Z domain and the *x′*-axis of C domain close to the minus *z*-axis of Z domain. It should be noted that the marginal distribution of the *z′*-axis is relatively broad but the marginal distribution of the *x′*-axis is narrow. Additionally, uncertainties in the observed RDC values are convolved into the CDIO and therefore no doubt artificially broaden it to some degree. Thus, the CDIO’s depicted in Figure 5 represent apparent CDIO’s. Accurate estimates of the experimental uncertainty contributions to the CDIO will require additional orthogonal alignments, but it is unlikely to represent more than 10% of the variance, based on the average RMSD values given in Figure S5b. In Figure 5c, the region bounded by the 60% probability threshold takes 9.7% of the entire interdomain orientational space and it has a probability of 40%. The shape of this joint distribution suggests that the flexible linker enables the two domains to sample a relatively large range of the interdomain orientational space but somewhat restricts the orientation of the C domain’s *x′*-axis in the vicinity of the Z domain’s *x*-*y* plane. It should be noted that the ribbon diagrams in Figure 5 represent the most probable interdomain orientations of each solution, but the solutions are not single structures. Rather, each solution is a distribution of structures depicted by the DoS figures. The DoS figures are meant to replace atomistic figures when presenting conformational distributions.

We have explored the impact of a single Normal CDIO assumption on the analysis of the experimental data by refitting the RDC residuals to an additional Normal mode (see Section 5 of Supplementary Information). In the case of the ZLBT-C data, this additional mode amounts to only 13% of the total probability and therefore contributes little to the overall distribution (Figure S9). However, in cases where multiple interdomain specific interactions occur, one might expect multiple modes to be significantly populated. In these cases, this extension, which we refer to as the Bingham mixture model, could be quite useful.

### 5.5 The energetically favorable CDIO of SpA

The thermodynamics of a binding reaction can be broken into two distinct parts: 1) the rearrangement of the binding partners’ conformational ensembles to those found in the complex and 2) desolvation of the binding interface and the formation of intermolecular interactions. The first part has been called “induced fit” [30] and occurs in nearly all biomolecular recognition events. The second part corresponds to the “lock and key” [31] portion of the binding reaction. Because the underlying molecular mechanisms associated with these two parts are distinctly different, structural-thermodynamic insights into them both are crucial to understanding and optimizing macromolecular recognition.

Figure 6 depicts this thermodynamic analysis of a receptor-ligand binding reaction, with a focus on the conformational rearrangement of the receptor, here assumed to be a molecule with two rigid domains linked by a short flexible linker. The conformational rearrangment treated here is an interdomain reorientation, ignoring translational rearrangements to which our RDC experiments are blind (see Section 9.7). Note that the induced fit step always involves an increase in receptor energy because if the binding competent conformational ensemble were the lowest energy state, it would exist even in the absence of ligand. For favorable binding reactions, this unfavorable conformational energy is more than compensated for by favorable interfacial energies when the complex forms. Note also that there is no mechanistic information in this analysis. That is, binding to the original conformational ensemble could precede conformational rearangement or *vice versa*. Only kinetic experiments could determine the sequence of events. However, as with any coupled reaction, thermodynamic parameters can be broken into components, whether the associated steps are real or hypothetical.

**Figure 6:**
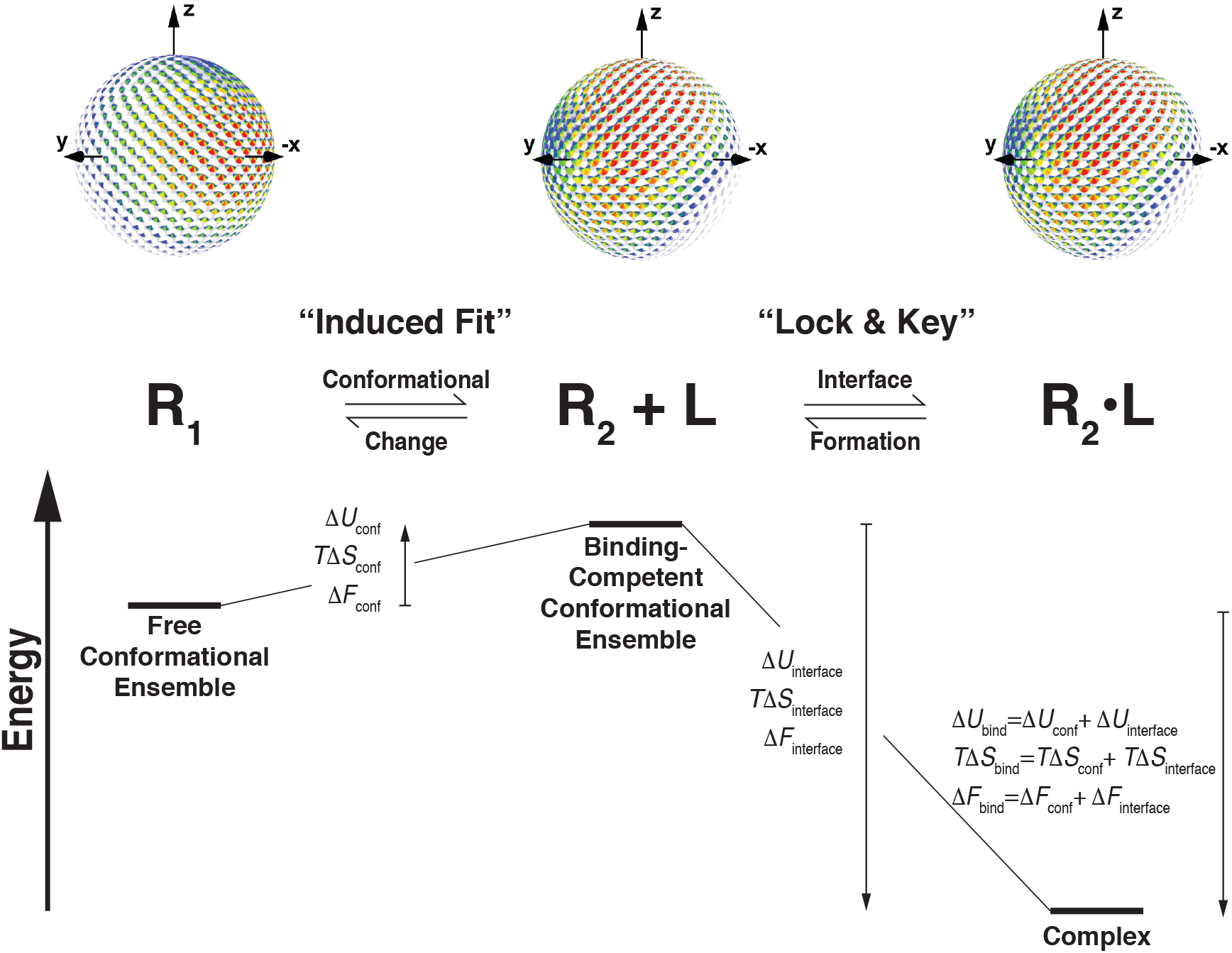
Thermodynamic analysis of the unfavorable reorientational contribution to the energy of molecular recognition. The binding reaction is broken into two parts: a change in the conformational ensemble of the two-domain receptor from R_1_ (the ensemble that exists in the absence of ligand) to R_2_ (the receptor ensemble when it is bound to ligand); and the desolvation and formation of the receptor-ligand interface. The order of events is arbitrary (see text) and the overall binding thermodynamics can be deconstructed without regard to mechanism. The CDIO of R_1_ is based on ZLBT-C RDC measurments and that of R_2_ is based on a simulation of the ZLBT-C/IgG complex (see text). Note that the R_2_ DoS figure is the same in R_2_ and R_2_·L. When the energetic contributions from the conformational change and interface formation reactions are added, they give the thermodynamics of the overall coupled binding reaction. As described in the text, the three thermodynamic parameters: ΔU_conf_, *ΔS*_conf_ and *ΔF*_conf_ can all be obtained from appropriate integration of the CDIO probability density functions R_1_ and R_2_ (see Section 9.7).

Because a CDIO is a valid Boltzmann distribution, our method allows the calculation of the difference in free energy, internal energy, and entropy between two CDIO’s (see Section 9.7). When these represent free and bound states of a receptor, the thermodynamic values obtained correspond to the receptor’s “induced fit” reorientation portion of the binding thermodynamics. In principle, if the same approach were applied to the ligand as well, then the free energy penalty of binding partner interdomain rearrangement could be accurately determined. It should be noted that intradomain rearrangements not captured by RDC’s would not be included in this calculation, but may be accessible to other structural methods.

The CDIO model allows a straightforward calculation of the difference in free energy, internal energy, and entropy between two distributions (see SI). This information can provide insights into the contribution of conformational reorientation to binding energetics. The thermodynamic parameters of the conformational change step and their contribution to the overall binding thermodynamics can be calculated using CDIO models of the free and bound protein. In the future, we plan to experimentally determine the CDIO for ZLBT-C bound to two IgG molecules. In place of this experimental data, we simulated the interdomain orientational distribution of ZLBT-C bound to IgG as a proof-of-principle. We modeled the complex as two IgG1 molecules bound to a Z-C model using the same interface observed in our previously determined C-Fc crystal structure [12]. The core F_ab_, F_c_, Z and C domains were treated as rigid bodies while the backbone conformations of the flexible 9-residue F_c_-F_ab_ linkers and the 6-residue Z-C linker were sampled using RanCh [28] to generate a 10,000 member ensemble, as described above.

Quantitative comparison of the simulated ZLBT-C/IgG_2_ CDIO to that of solution 2 (Figure S8) gives the re-orientational contributions to the Helmholtz free energy of binding (*ΔF*) of 0.8 kcal/mol, internal energy of binding (*ΔU*) of 0.7 kcal/mol and entropy (*ΔS*) of 0.33 cal/(mol·K). It should be noted that because a significant change in compressibility upon re-orientation is unlikely, Helmholtz free energy is approximately equal to Gibbs free energy (*ΔF* ≈ *ΔG*) and internal energy is approximately equal to enthalpy (*ΔU* ≈ *ΔH*) in this case. To compare these values to those of other possible CDIOs, we systematically sampled *SO*(3) space with 331,776 different CDIOs whose variances were each set equal to those of the simulated ZLBT-C/IgG_2_ CDIO. We then calculated the *ΔF* and *ΔU* of each CDIO relative to the experimentally determined ZLBT-C CDIO. The *ΔF* of solution 2 is in the 23^*rd*^ percentile of all values, which range from 0.3 to 1.8 kcal/mol. Note that the reorientational penalty is not 0 even when the mean of the sampled CDIO matches that of the free ZLBT-C CDIO because the variance of the ZLBT-C/IgG_2_ CDIO is predicted to be lower than that of the observed free ZLTB-C CDIO. The large difference between the energy cost of 0.8 kcal/mol and the maximum cost of 1.8 kcal/mol indicates that the maximum probability orientation of solution 2 is posed in a favorable orientation to bind antibodies. Altering the maximum probability orientation could significantly increase the conformational energy cost of binding, which suggests that the observed interdomain orientational distribution has been evolutionarily optimized.

## 6 Discussion

Previous studies have shown that calculations in structural biology that rely on discrete conformational ensembles are brittle. Instead, the continuous nature of proteins must be taken into account in order to avoid overtting and brittle changes in energetics as one moves between discrete structures. Only in the limit can a discrete ensemble approximate a continuous distribution adequately, and this limiting case is far too expensive for any computer. In the absence of a continuous model, current protein design methods start with one or several discrete structures [25, 26, 27]. The methods then model conformational distributions by using either molecular dynamics or by sampling and weighting nearby conformations using an empirical molecular mechanics energy function. A continuous model can certainly help this scenario by directly using experimentally determined flexibility for protein design.

The broad, albeit highly anisotropic, orientational distribution of ZLBT-C indicates that the molecule samples a large region of interdomain orientational space, as a result of high structural flexibility. As previously noted, this structural flexibility may be crucial to the large functional plasticity displayed by this important *S. aureus* virulence factor [13]. The interdomain orientational distribution is certainly influenced, if not dominated, by the six-residue flexible linker. The linker sequences are nearly identical in the 4 linkers in SpA-N and among more than 50 *S. aureus* isolates (unpublished observations). One possible reason for this unusual conservation is to maintain the same interdomain orientational distribution. If this hypothesis is correct, the interdomain orientational distribution is selected by evolutionary pressure as a functional phenotype of the protein. Future studies using a polyglycine linker of the same length will be used to test this hypothesis.

Based on the relative magnitudes of RDCs in the first and second domains, the ZLBT-C construct has a more flexible interdomain linker than two previously studied di-domain systems [1, 2]. Surprisingly, the region bounded by blue color in Figure 5c covers only 35% of the entire 3D interdomain orientational space but it has a total probability of 80%. Although the distribution is broad, it certainly shows a strong preference over a limited volume. Our results suggest that a binary classification of biopolymers as ‘flexible’ or ‘inflexible’ is an over-simplification. For flexible systems with different parts working jointly, quantifying the coupled conformational distribution is necessary to reveal the molecular mechanism underlying concerted motions and the relationship between motion and function. Based on the continuous conformational distribution, we can calculate and separate the contributions of enthalpy and entropy to binding very easily, by computing a simple integral. Having calculated the contributions of enthalpy and entropy to binding for the ZLBT-C di-domain system (see Section 9.7), we now believe the interdomain orientation distribution may have been evolutionarily optimized to reduce the conformational energy cost of binding.

Finally, the CDIO model we present here offers an example of a starkly different approach to interpreting RDC observations in flexible di-domain systems. By avoiding pre-enumeration of atomistic ensemble models, the CDIO approach simplifies the process of generating a probabilistic structural model and makes it easier to interpret. The DoS representation makes interdomain orientational probability distributions easier to visualize and relate to a conventional atomistic representation using a single set of Cartesian coordinates. By providing the algorithms and software necessary to fit a CDIO model and display it as a DoS figure, we hope our approach will be easy to adopt.

## 7 Acknowledgments

We thank Hashim Al-Hashimi, Mark Hallen, Swati Jain, Pablo Gainza, James Prestegard, Ken Dill, Hunter Nisonoff and Rachel Kositsky for their helpful discussions. A portion of this work was conducted at Duke NMR center. We would like to thank Dr. Ronald Venters, Dr. Anthony Ribeiro and Donald Mika for their constant help. The work was supported by NIH grant R01-GM-78031 (to B.R.D.), R01-GM-118543 (to B.R.D.) and NIH grant R01-GM-081666 (to T.G.O.).

## 8 Author Contributions

Y.Q., A.W.B., B.R.D. and T.G.O. designed the experiments. Y.Q. performed the experiments. Y.Q., J.W.M., A.Y., F.T. and B.R.D. designed the algorithm. Y.Q. and F.T. implemented the algorithm. Y.Q., J.W.M., F.T., A.W.B., B.R.D. and T.G.O. analyzed the data. Y.Q., A.W.B., B.R.D. and T.G.O. prepared the figures and wrote the paper.

## 9 Materials and Methods

### 9.1 Molecular Cloning and Protein Preparation

The 5B and ZLBT-C genes were synthesized by GENEWIZ, Inc. and then cloned into pAED4. The design of ZLBT-C was based on the Z-L2LBT construct [19]. The amino acid sequence of ZLBT-C is ‘MVDNKFNKEQQNAFYEILHLPNLNEEQRNAFIQS LKDYIDTNNDGAYEGDELQSANLLAEAKKLNDAQAPKADNKFNKEQQNAFYEILHLPNLTEEQRNGFIQSLKDDPSVSKEILAEAKKLNDAQAPK’. The NHis-ZLBT-C gene (ZLBT-C with a 6×His tag fused to the N-terminus) was constructed by inserting the ZLBT-C into pET28a (Novagen) and removing the thrombin cleavage site. The amino acid sequence of NHis-ZLBT-C is ‘MGSSHHHHHHSSGLVPRGSH MVDNKFNKEQQNAFYEILHLPNLNEEQRNAFIQSLKDYIDTNNDGAYEGDELQSANLLAEAKKLNDAQAPKADNKFNKEQQNAFYEILHLPNLTEEQRNGFIQSLKDDPSVSKEILAEAKKLNDAQAPK’. All plasmids were transformed into *E. coli* BL21(DE3) for expression. Non-labeled proteins were expressed at 37°C for 5 hrs in LB medium and isotope-labeled proteins were expressed in M9 medium. Proteins were purified first by acid precipitation with 5% (v/v) HAc solution, then by a cation-exchanging SP Sepharose (GE Healthcare) column or a Ni-NTA affinity column. The purity of all products was confirmed by SDS-PAGE and their masses were confirmed with electro-spray ionization mass spectroscopy.

### 9.2 Determination of the Dissociation Constant between [*Tb*^3+^] and LBT constructs

TbCl_3_ was titrated into 0.5 *μ*M protein solution (ZLBT-C or NHis-ZLBT-C) to final concentrations ranging from 0.25 to 10 μM. The protein solution is in the buffer of 25 mM MOPS (pH7.2) and 100 mM KCl. The tyrosine residue in the middle of LBT was excited at 280 nm and the emission of the Tb^3+^ ion was observed at 495 and 545 nm. The emission intensities were fit to the following binding equation to obtain the dissociation constant *K_D_*:

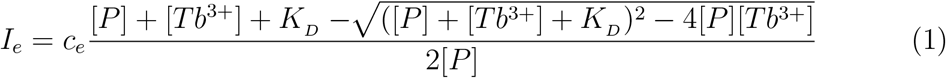

where *I_e_* is the emission intensity, *c_e_* is the emission coefficient, [*P*] is the protein concentration and [*Tb*^3+^] is the concentration of Tb^3+^ ions.

### 9.3 NMR Measurements and Analysis

NMR spectroscopy was performed on Varian instruments (14.1T and 18.8T) with a Brüker console (14.1T) and an INOVA console (18.8T). 5B NMR samples were prepared with 1mM protein in 50 mM NaAc/HAc (pH 5.5) and 100 mM NaCl with 10% D_2_O. ZLBT-C and NHis-ZLBT-C samples were prepared with 250-500 *μ*M protein in 25 mM MOPS (pH 7.2) and 100 mM KCl with 10% D_2_O. The ^15^N-HSQC spectra of 5B and ZLBT-C were sequentially assigned using two 3D NMR experiments, HN-CACB and CBCA(CO)NH. The assignments of NHis-ZLBT-C were transferred from the assignments of ZLBT-C. For 5B, all experiments were performed at 30°C. ^15^N T1, T2 measurements and ^15^N Heteronuclear NOE experiments were conducted and spectra were processed by NMRPipe. R1, R2 rates and NOE ratios were calculated using the program NMRViewJ. The spectral density function evaluated at three frequencies, 0, *ω*_N_ and 0.87*ω*_H_ Hz, were calculated from R1, R2 rates and NOE ratios and were subsequently used to generate the Lipari-Szabo map [16]. Order parameters were also derived from the three spectral density function values using an anisotropic tumbling model. For ZLBT-C and NHis-ZLBT-C, all experiments were performed at 25°C. ^15^N HSQC IPAP experiments [20] were conducted with 1024 complex points in the direct dimension and 128 points in the indirect dimension. The samples for these experiments contain 1:1.1 ratio of protein and LuCl_3_/DyCl_3_/TbCl_3_. The sample with LuCl_3_ was used to measure the J coupling and serves as a reference. The spectra were also processed by NMRPipe. The center of each peak was measured by fitting the peak to a mixture model with both Gaussian and Lorentzian components. The fitting program was written in Mathematica. RDCs were measured by calculating the difference between the coupling constants in the presence of diamagnetic ion Lu^3+^ and paramagnetic ions Dy^3+^/Tb^3+^. The Saupe tensor of each domain in each alignment was calculated by the SVD method [21]. The Z domain structure (PDB code: 1Q2N) and C domain structure (PDB code: 4NPE) were used for the calculation. The Q factor associated with each dataset reflects the level of noise both in the RDC data and the input structure. The noise level is generally considered low when the Q factor is below 0.3 [22]. The Q factor is calculated as:

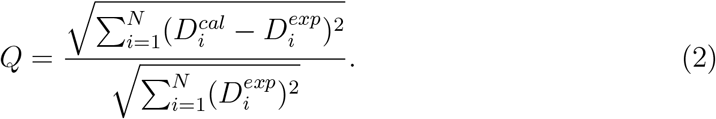

The fitted Saupe tensors were used to generate orthogonal alignments following the OLC method [22]. Programs for Saupe tensor calculation and the OLC method were also written in Mathematica.

### 9.4 Interdomain Motion Modeling and Data Fitting

The interdomain motions were modeled as a Bingham distribution on *SO*(3) [23]. The Bingham model on *SO*(3) is a unimodal distribution, which takes the following form:

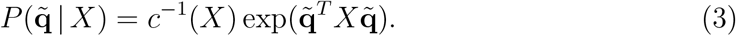

In Eq. (3), *c*^−1^(*X*) is the normalization factor. *q* ∈ *SO*(3) is a rotation and 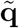 is the 4D unit quaternion representation of *q*. Unit quaternions can represent interdomain orientations. A unit quaternion can be described by four parameters. The four parameters represent a single-axis rotation, or the resulted orientation. Three of the parameters represent a 3D unit vector, which is the rotation axis. The fourth parameter describes a rotation angle around the axis. The unit quaternion representation of orientation preserves Harr measure and avoids singularity [29]. Matrix *X* encodes the mean and variances of the Bingham model on *SO(3)*. It is a symmetric 4 × 4 matrix with a constant trace which thus has 9 degrees of freedom (DOFs).

The relationship between the Saupe tensor of Z domain and the Saupe tensor of C domain in the same alignment is described by a 5 × 5 transformation matrix Q with 25 real elements, as shown in the following equation:

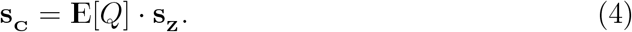

In Eq. (4), **s_C_** and **s_Z_** are vectorized Saupe tensors with 5 elements each (Supplementary Information). The *Q* matrix is a function of the interdomain orientation, *R* ∈ *SO*(3). **E**[*Q*] is the expectation value of the Q matrix over the rotation space *SO*(3) given a probability distribution for *R*. Using the Bingham model, **E**[*Q*] becomes a function of *X*. Based on Eq. (4), the objective function used in the fitting algorithm is defined as:

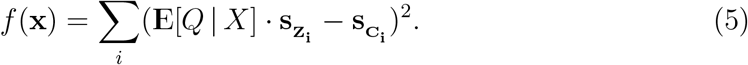

In Eq. (5), *X* contains the array of parameters to define the Bingham distribution; **E**[*Q* | *X*] is the calculated expectation value of the *Q* matrix given the parameters *X*; *s_Z_i__* is the *i* ^th^ experimental alignment tensor of Z domain; and *s_C_i__*, is the *i* ^th^ experimental alignment tensor of C domain. The derivation and the elements of the *Q* matrix are given in the Supporting Information.

The fitting algorithm is a branch-and-bound algorithm that exploits a decomposition of the *Q* matrix when assuming the Bingham model. The *Q* matrix was decomposed into the product of three matrices. The DOFs of the three matrices constitute spaces *S*^3^, 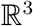 and *S*^3^, respectively. *S*^3^ is the unit 3-sphere in 4-dimensional space 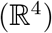. To distinguish the two rotation spaces, the space corresponding to a left rotation is notated as 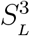 and the one corresponding to a right rotation is notated as 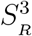. The fitting algorithm has three steps. In the first step, the space 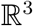 was divided into subregions, which were subsequently bounded and pruned. In the second step, the remaining 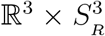 space was branched, bounded and pruned. The final step of algorithm exhaustively enumerated the remaining space in 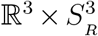 after the two-step pruning. Potential solutions were scored based on the objective function defined in Eq. (5). Solutions with the best scores were returned. A detailed description of the fitting algorithm is provided in the Supplementary Information. The CDIO fitting program and the DoS rendering Mathematica notebook are freely available, as open-source software (http://www.cs.duke.edu/donaldlab/software/cdio.php). This makes our code and algorithm generally available to the community.

### 9.5 Calculation of Clash Scores for Distributions

Given a low probability region defined by a simulation, the probability of the region is defined as the floor probability *p_f_*. The floor probability represents the reasonable amount of probability existing in the region. In addition, define a ceil probability *p_c_* as the largest probability drawn from an equal volume region in the simulated distribution. The clash score for a distribution is calculated as following:

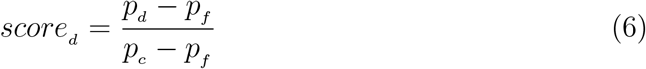

where *p_d_* is the probability of a distribution falling within the low probability region.

### 9.6 Simulation of the ZLBT-C/IgG_2_ complex

Three structure models, Z domain (PDB code: 1Q2N), C domain-F_c_ complex (PDB code: 4WWI) and a full antibody molecule (PDB code: 1IGT), were used in the simulation generated by RanCh [28]. After generating a 10,000 member ensemble, we converted the discrete ensemble into a continuous distribution by assigning uniform weights to each conformer and using an isotropic Bingham kernel (See the convolution module in the CDIO software package).

### 9.7 Calculation of Thermodynamic Parameters from Distributions

When a molecule (referred to hereafter as the receptor) binds its binding partner (the ligand), its conformational distribution changes. Given the conformational probability density function (PDF) *P*_1_(*x*) in the absence of ligand and the PDF *P*_2_(*x*) in the presence of the ligand, thermodynamic parameters (*ΔF*, *ΔU*, and *ΔS*) for reorientation of the receptor from the free to bound ensembles can be calculated assuming that a probability in the reference distribution *P*_1_(*x*) and its corresponding state energy *E*_1_(*x*) satisfies the Boltzmann distribution relationship:

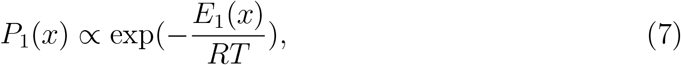

where *x* represents the three orientational degrees of freedom. Note that conformational changes leading to interdomain translation are not considered because the RDC measurements are blind to such changes. However, other experiments (e.g., FRET or pseudocontact shifts) might provide such complementary information. Here, we only focus on the receptor conformational change contributions to the binding thermodynamics. We define the receptor system as an ensemble of conformations adopted by the unliganded receptor. The internal energy of the receptor ensemble before binding is

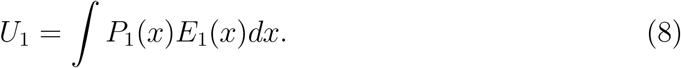

The conformational internal energy of the receptor ensemble after binding is

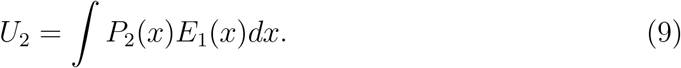

So the change in the internal energy of the receptor ensemble due to the binding reaction is

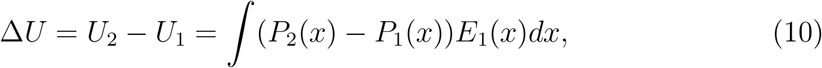

because the Boltzmann equation is:

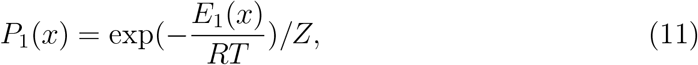

where *Z* is the partition function for the unbound receptor ensemble, we have:

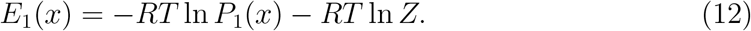

By substituting Eq. (12) into Eq. (10), we have:

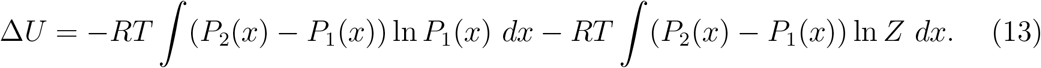

The second term on the right-hand side in Eq. (13) is zero because both PDF’s *P*_1_(*x*) and *P*_2_(*x*) are normalized and their integrals equal 1. Consequently, we have:

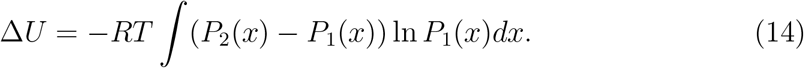

In addition, *ΔS* can be calculated using the probability form of the Boltzmann equation:

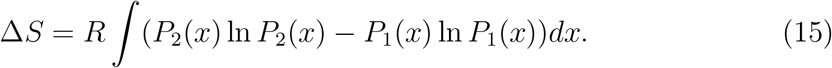

Consequently, the Helmholtz free energy can be calculated from *ΔU* and *ΔS* as:

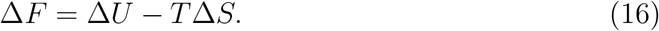

We used the above equations to calculate *ΔF*, *ΔU* and *ΔS* from our computed CDIO distributions.

### 9.8 Disk-on-sphere representation

The challenge of visualizing orientation probability distributions is that they are joint in the three DOFs of *SO*(3). Displaying the correlation between the DOFs requires a 4-dimensional representation. Each interdomain orientation can be specified by the direction **e**_*z*_ of the second domain’s *z*-axis and the direction **e**_*x*_ of the second domain’s *x*-axis. In the disk-on-sphere representation, location of the center of a disk on a 2-sphere corresponds to **e**_*z*_, while the direction of a radial line on the disk corresponds to **e**_*x*_. The radial lines are color coded to represent their associated probabilities. The transformation from a quaternion to its corresponding vectors, **e**_*z*_ and **e**_*x*_, is achieved by:

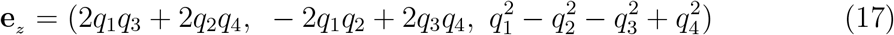

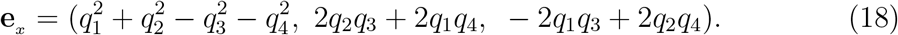

The disk-on-sphere representation preserves the features of the rotation space *SO*(3) better than an Euler angle representation. The distribution of **e**_*z*_ can be plotted on a spherical surface without distortion. In our case to draw it, the distribution of **e**_*z*_ is discretized on the sphere to incorporate disks representing the conditional distribution *p*(**e**_*x*_ | **e**_*z*_), the disk-on-sphere representation expresses most features of not only the rotation space *SO*(3) but also distributions on *SO*(3). By zooming in, a finite discretization of **e**_*z*_ can be calculated and drawn from the underlying continuous distribution at any desired resolution.

## Footnotes

1 RDC: : residual dipolar coupling; DoS: : disk-on-sphere; SpA: : Staphylococcal protein A; SpA-N: : N-terminal half of SpA; LBT: : lanthanide binding tag; CDIO: : continuous distribution of interdomain orientations;

